# Pleiotropy or linkage? Their relative contributions to the genetic correlation of quantitative traits and detection by multi-trait GWA studies

**DOI:** 10.1101/656413

**Authors:** Jobran Chebib, Frédéric Guillaume

## Abstract

Genetic correlations between traits may cause correlated responses to selection. Previous models described the conditions under which genetic correlations are expected to be maintained. Selection, mutation and migration are all proposed to affect genetic correlations, regardless of whether the underlying genetic architecture consists of pleiotropic or tightly-linked loci affecting the traits. Here, we investigate the conditions under which pleiotropy and linkage have differential effects on the genetic correlations between traits by explicitly modeling multiple genetic architectures to look at the effects of selection strength, degree of correlational selection, mutation rate, mutational variance, recombination rate, and migration rate. We show that at mutation-selection(-migration) balance, mutation rates differentially affect the equilibrium levels of genetic correlation when architectures are composed of pairs of physically linked loci compared to architectures of pleiotropic loci. Even when there is perfect linkage (no recombination within pairs of linked loci), a lower genetic correlation is maintained than with pleiotropy, with a lower mutation rate leading to a larger decrease. These results imply that the detection of causal loci in multi-trait association studies will be affected by the type of underlying architectures, whereby pleiotropic variants are more likely to be underlying multiple detected associations. We also confirm that tighter linkage between non-pleiotropic causal loci maintains higher genetic correlations at the traits and leads to a greater proportion of false positives in association analyses.

## Introduction

Both pleiotropy and linkage disequilibrium create genetic correlations between traits so that traits do not vary independently of one another (Wright, 1977; Arnold, 1992; Walsh and Blows, 2009). Under natural selection, this process can prevent a combination of traits from reaching their respective optimum trait values favored by natural selection (Falconer and Mackay, 1996). Likewise, under artificial selection it can constrain breeders from improving one trait due to undesired changes in another, and in medical gene targeted therapy treatments it can cause adverse side-effects (Wright, 1977; Parkes et al., 2013; Visscher et al., 2017; Wei and Nielsen, 2019). Pleiotropy may cause genetic correlation because one gene’s product (e.g., an enzyme or a transcription factor) has more than one target and therefore affects more than one trait or because one gene’s product belongs to a metabolic pathway that has more than one downstream effect (Hodgkin, 1998; Stearns, 2010; Wagner and Zhang, 2011). Linkage disequilibrium (LD) may be the result of a set of loci in close physical proximity on a chromosome that makes a set of alleles at those loci less likely to be split up by recombination and therefore more likely to get passed on together from one generation to the next. But other mechanisms leading to the transmission of one combination of alleles at separate loci over another combination, can also generate LD and create genetic correlations between traits that those loci affect (e.g., assortative mating, environmental correlations) (Falconer and Mackay, 1996).

One of the main objectives of a genome-wide association study (GWAS) is to identify causal genetic variants underlying one or more traits. GWASes leverage the rapid increase in genomic sequencing to find correlations between traits and genotypes, and their success is dependent on the effect sizes of the loci and the distinction between phenotypes. GWASes have had success in associating genetic variants with traits of interest, which have allowed researchers to find the molecular underpinnings of trait change (Visscher et al., 2017). Moving from one trait to two or more trait associations can lead to discovering pleiotropic loci (Saltz et al., 2017). One GWAS using 1094 traits and 14,459 genes, found that 44% of genes were “pleiotropic”, but this was determined by assigning genetic variants to the closest gene and even to both flanking genes when the genetic variant was intergenic (Chesmore et al., 2018). This conflates linkage and pleiotropy, and the chain of causality (Platt et al., 2010). Another study, found 90% of genes and 32.4% of SNPs were associated with more than one trait domain, but they could not rule out SNPs associated with traits due to linkage disequilibrium (Watanabe et al., 2018). Unfortunately, determining whether genetic variant associations and trait correlations are actually the result of pleiotropy or linkage is difficult since they often map to large regions of genomes, or are in intergenic regions and don’t associate with the closest genes (Flint and Mackay, 2009; Zhu et al., 2016; Peichel and Marques, 2017; Visscher et al., 2017). Distinguishing between the two types of genetic architectures is important for understanding the underlying molecular functions of the traits, and determining how the traits may be deferentially affected by selection (Lynch et al., 1998; Barrett and Hoekstra, 2011; Saltz et al., 2017). This is salient at a time when an increasing number of traits of interest (e.g., human diseases) appear to be affected by loci that affect other traits, and especially when targeted gene therapy clinical trials are more widespread than ever (Edelstein et al., 2007; Cai et al., 2016; Pickrell et al., 2016; Visscher and Yang, 2016; Chesmore et al., 2018; Ginn et al., 2018). There are potentially negative implications for gene therapy because fixing a gene underlying one disease might increase risk for another disease. For example, some genetic variants that are associated with greater risk of Ankylosing spondylitis are also associated with less risk of Rheumatoid arthritis, and so “fixing” one gene would have undesired sideeffects in this case (Parkes et al., 2013; Gratten and Visscher, 2016).

But the evolutionary dynamics of pleiotropic versus linked loci in creating genetic correlations are expected to be different, since pleiotropy requires only one mutation to affect multiple traits and build-up genetic correlations, and linked pairs require two. Mutation rate should be an important factor distinguishing pleiotropy and linked pairs because single mutations affecting more than one trait provides the opportunity for combinations of effects to match patterns of correlational selection better than linked loci that affect one trait at a time. Thus, linked pairs may require high mutation rates to maintain genetic correlations. Recombination can also reduce genetic correlations between traits by breaking up associations between alleles at linked loci, but the same cannot occur with a pleiotropic locus (but see Wagner et al. (2007) for other mechanisms to alleviate pleiotropic constraints). Polygenic analytical models attempting to approximate the level of genetic variance and covariance at mutation-selection balance in a population suggest that tight linkage between pairs of loci affecting separate traits “is nearly equivalent to” pleiotropic loci affecting both traits (Lande, 1984). Therefore, genetic correlations between traits can be approximated using previously elucidated pleiotropic models under certain conditions (Lande, 1980, 1984; Turelli, 1985). On the other hand, more recent extensions of Fisher’s Geometric Model (Fisher, 1930) predict that pleiotropic mutations, compared to mutations that affect only one trait, are less likely to be beneficial overall since a beneficial effect on one trait may be detrimental to others (Orr, 1998; Otto, 2004). The detrimental effect of pleiotropy is exacerbated when increasing the strength of selection or with very strong correlational selection between traits, since both reduce the amount of phenotypic space where mutations are beneficial (unless pleiotropic effects are aligned with the fitness surface created by correlational selection). This detriment is not present for linked loci affecting separate traits since their beneficial mutations will not have the collateral effects of pleiotropy. These, therefore suggest that linkage and pleiotropy may have differential effects on genetic variance and covariances depending on mutation, recombination and selection regimes, but this comparison was not fully explored in any previous model.

Lande (1984) predicted that when loci affecting *different* traits are tightly linked, and there is strong correlational selection between traits, recombination rates between loci affecting different traits can strongly affect genetic correlations between traits, when selection is weak and mutation rates are relatively high. In an extreme case where there is complete linkage between pairs of loci affecting different traits (the recombination rate is 0), and no linkage between sets of these pairs of linked loci (the recombination rate is 0.5), then he determined that the maximum genetic correlation due to linkage may be almost as large as the extent of correlational selection, which can be calculated from the (per linkage group) genetic covariance between traits and the genetic variances, respectively, as:

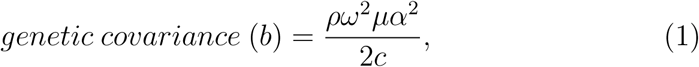

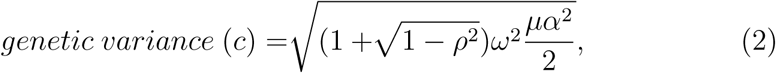

where *ρ* is the extent of correlational selection acting between the traits, *ω*^2^ is the strength of selection (with lower values representing stronger selection), *µ* is the per-locus mutation rate, and *α*^2^ is the per-locus mutation variance. If there is equal variances among traits then the genetic correlation is calculated as:

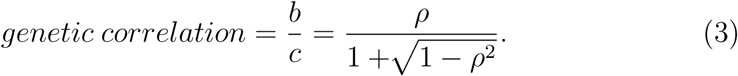

From these equations we see that, even in the absence of pleiotropy, genetic covariance may arise from linkage disequilibrium, and depends on both the strength of correlational selection between traits and selection on each trait, as well as on the mutational inputs (mutation rates and mutational variances) of the genes affecting those traits. Yet, from equation (3), the resulting genetic correlation among traits is independent of the genetic architecture of the traits. Lande goes on further to state that the case of complete linkage between pairs of loci affecting different traits is “equivalent to a lesser number of loci with pleiotropic effects”, but this is not quantified nor is the scaling of the two examined. We seek to quantify the equivalence of pleiotropy and linkage in their ability to maintain equilibrium levels of genetic (co)variation under the same conditions. We also wish to extend this to look at a range of linkage distances, selection variances and correlations, and mutation rates and variances, to look at the relative effects of each.

The expectations given by Lande are only expected to be accurate under conditions where mutation rates are high compared to the strength of selection on the traits of interest (Turelli, 1984; Turelli and Barton, 1990). When mutation rates are lower (< 10^−4^), predictions for equilibrium levels of genetic variation break down and are better approximated by the “house-of-cards” model (Kingman, 1978; Turelli, 1984). Analytic predictions for equilibrium levels of genetic covariation between traits due to linkage disequilibrium, on the other hand, have not been well explored for the “house-of-cards” model (Bürger, 2000).

Additionally, levels of trait genetic covariation can be influenced by other evolutionary processes that affect allele frequencies, and the covariation of allelic values in a population (e.g., migration (Guillaume and Whitlock, 2007), drift (Griswold et al., 2007), inbreeding (Lande, 1984), and phenotypic plasticity (Draghi and Whitlock, 2012)). Migration affects genetic covariation because when it is sufficiently high (relative to selection in the focal population), then combinations of alleles coming from a source population will also be maintained in the focal population. This can lead to higher genetic covariation between traits in the focal populations, whether the combinations of alleles immigrating are (more likely to be) correlated in their effects on those traits or not (Guillaume and Whitlock, 2007). Migration may also have different effects depending on whether the genetic architecture is pleiotropic or made up of linked loci, but this has not been explored.

Here, we are interested in the conditions in which pleiotropic architectures behave similarly or differently to architectures with tight physical linkage between loci affecting different traits, with respect to their effects on genetic correlations between the traits. We use computer simulations to investigate whether the effect of evolutionary forces on the genetic correlation between traits is dependent on the type of genetic architecture, and how. We focus on the relative contributions of selection, mutation and migration to the build up of genetic correlation between traits having different genetic architectures. We show that unless mutation rates are high, genetic architectures with tight linkage between loci maintain much lower equilibrium genetic correlations than pleiotropic architectures. Even when mutation rates are high, other evolutionary forces affecting equilibrium levels of genetic correlation still show a difference between architectures but to a much lesser extent. Additionally, we simulate genomic single-nucleotide polymorphism (SNP) data sets using the different architectures, and show that map distances between causative and non-causative QTL affect false positive proportions in GWA analyses.

## Materials and Methods

We modeled four different genetic architectures in a modified version of the individual-based, forward-in-time, population genetics simulation software Nemo (Guillaume and Rougemont, 2006; Chebib and Guillaume, 2017). Nemo was modified to allow single non-pleiotropic loci to affect different quantitative traits. To compare how pleiotropy and linkage differentially affect the genetic correlation between traits, we modeled a set of 120 pairs of linked, non-pleiotropic loci, and a set of 120 pleiotropic loci affecting the two traits. We varied the recombination distance between the two non-pleiotropic loci of each pair with distances 0cM, 0.1cM, or 1cM (Figure 1). Pairs were unlinked to other pairs. The pleiotropic loci were also unlinked to each other. The recombination rates chosen represent no recombination between linked loci, as well as an average and an extreme value of recombination at “hotspots” in the human genome, respectively (Myers et al., 2006). All loci had additive effects on the traits.

**Figure 1:**
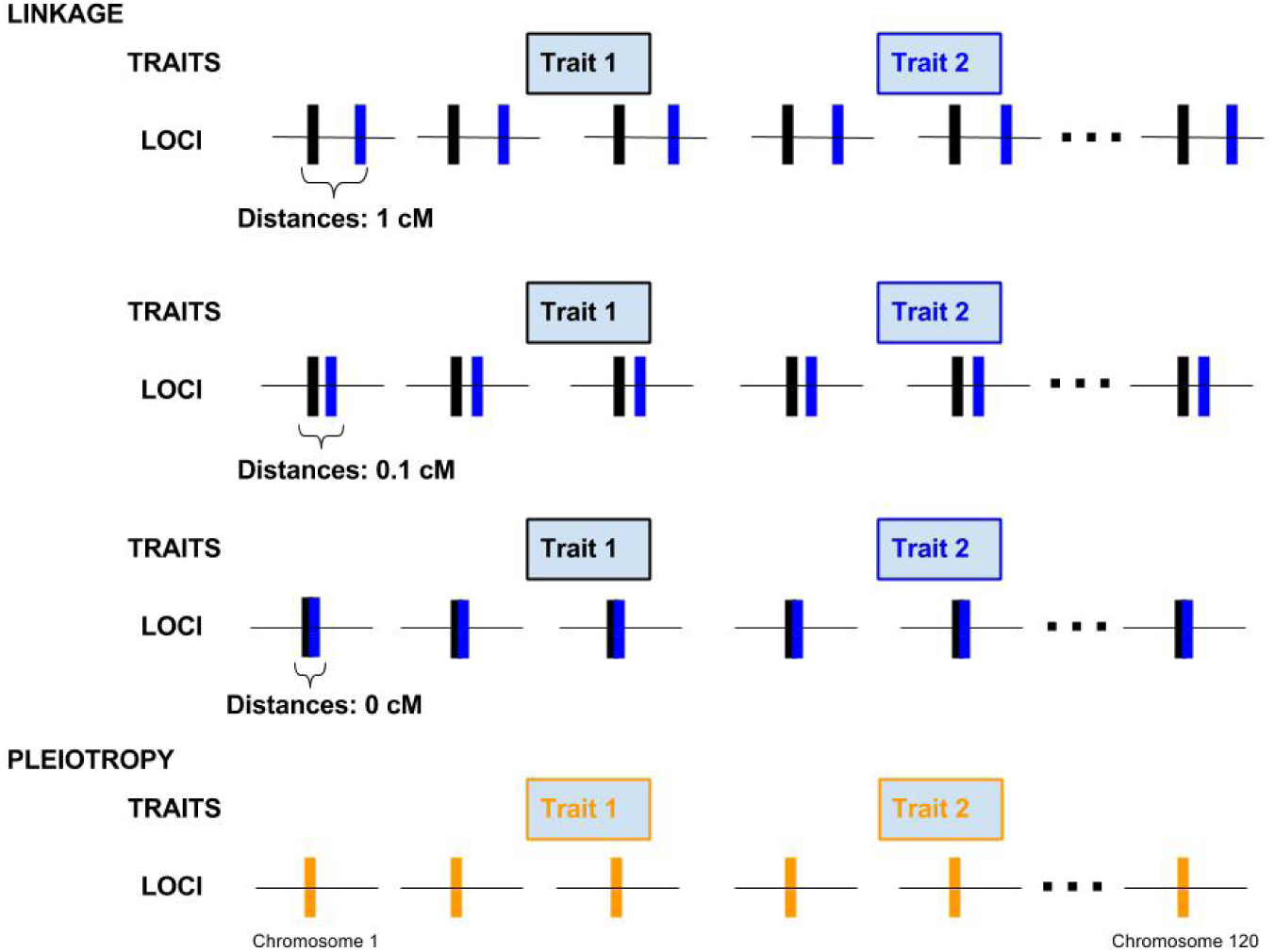
Four genetic architectures showing the distribution of loci on 120 chromosomes. In the case of linkage architectures, pairs of loci affecting the two different traits on each chromosome are either 1, 0.1 or 0 cM apart. In the case of the pleiotropic architecture, each locus on each chromosome affects both traits.

Unless otherwise specified, each simulation was run with 5,000 initially monomorphic (variation is gradually introduced through mutations), diploid individuals for 10,000 generations achieving mutation-selection(-migration) balance in order to observe general patterns of genetic correlation in the near-absence of drift. Individuals were hermaphrodites mating at random within a population, with non-overlapping generations. Phenotypes were calculated for each of the two traits modeled by summing the allelic values of all loci affecting one trait. Gaussian stabilizing selection was applied and determined the survival probability of juveniles, whose fitness was calculated as 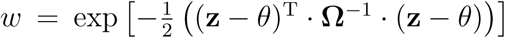, where **z** is the individual phenotype vector (initialized to the optimum values), *θ* is the vector of local optimal trait values (set to 10 for both traits in the focal population), and **Ω** is the selection variance-covariance matrix (*n × n*, for *n* traits) describing the multivariate Gaussian selection surface. To examine the effects of the strength of stabilizing selection on each trait and strength of correlational selection between traits, different sets of simulations were run with the diagonal elements of the **Ω** matrix set as *ω*^2^ = 50, or 100 (selection strength), and off-diagonal set to *ω*^2^ *× ρ_ω_* (where the correlational selection, *ρ_ω_* = 0.5 or 0.9). The strength of selection scales inversely with *ω*^2^ where a value of 100 corresponds to weak (but non-trivial) selection as opposed to correlational selection, *ρ_ω_*, where a value of 0.9 corresponds to strong correlational selection between traits (Lande, 1984; Turelli, 1984).

To examine the effects of mutational input on genetic correlation between traits, different sets of simulations were run with mutation rates (*µ*) of 0.001, 0.0001, or 0.00001, and moderate mutational effect sizes (*α*^2^) of 0.1, 0.01, or 0.001 (Turelli, 1984). Mutational effects at each non-pleiotropic locus were drawn from a univariate normal distribution (with a mean of zero) or a bivariate normal distribution (with means of zero and a covariance of 0) for pleiotropic loci. Mutational effects were then added to the existing allelic values (continuum-of-alleles model; Crow and Kimura, 1964). All loci were assumed to have equal mutational variance. No environmental effects on the traits were included.

To examine the effects of migration from a source population on genetic correlation between traits, additional sets of simulations were run with unidirectional migration from a second population (as in an island-mainland model with each population consisting of 5000 individuals) with backward migration rates (*m*) of 0.1, 0.01, and 0.001. The backward migration rate represents the average proportion of new individuals in the focal population whose parent is from the source population. The local optimum values for the two traits in the source population were set at 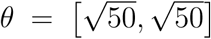 (10 units distance from the focal population’s local optimum). Both focal and source populations had weak stabilizing selection with a strength of *ω*^2^ = 100, the focal population had no correlational selection between the two traits and the source population had a correlational selection of *ρ_ω_* = 0 or 0.9. Fifty replicate simulations were run for each set of parameter values and statistics were averaged over replicates. Averages were also compared against analytical expectations laid out by Lande (1984) and reproduced here in Equations 1–3.

### Effects of genetic architecture on false positive/negative proportions in association studies

In order to elucidate the differential effects of pleiotropy and linkage on the detection of true causal genetic variants in association studies, a genome-wide association (GWA) analysis was performed on data simulated as described above (with only a single population), except that diallelic loci were used instead of a continuum-of-alleles model to better represent SNPs. Correlational selection values were chosen that provided equal on-average genetic correlations between traits for all genetic architectures of 0.2, 0.3, and 0.4, values frequently observed in both morphological and life-history traits (Roff, 1996). In the association study, a per-locus regression of trait values was performed over genotypes, and the (negative log 10) p-values of regression slopes were plotted with a Benjamini-Hochberg False Discovery Rate (FDR) cutoff to adjust significance levels for multiple tests (Benjamini and Hochberg, 1995). From this, we observed the number (and proportion) of false positives (linked loci that had no effect on a trait but whose regression slope p-values were above the FDR cutoff for that same trait) and false negatives (pleiotropic loci that had an effect on both traits but whose regression slope p-values were below the FDR cutoff for either trait). No correction for population stratification was performed during this analysis because each simulation had a single, large, randomly breeding population. Linkage disequilibrium values of *D′* and *R*^2^ between pairs of linked traits were also calculated using the **R** package genetics (v1.3.8.1) (Warnes et al., 2013). Statistics for number and proportion of false positives and negatives were obtained from the average over 20 replicate simulations of each genetic architecture. We also assessed the false positive rate on an additional set of neutral QTL linked to the causal loci. We simulated a set of 120 independent linkage groups with 200 neutral di-allelic QTL per group, evenly distributed on both sides of the central position occupied by the two causal QTL. Each linkage group was 1 cM long. The minimum recombination rate between two adjacent loci was 10^−5^. The neutral QTL were set in 10 successive windows of 0.05cM (∼50kb) on each side of the causal QTL. The two causal QTL were perfectly linked (0cM) and non-pleiotropic. The simulations were run for 50,000 generations and 10 replicates.

## Results

### Effects of genetic architecture on genetic correlation at mutation-selection balance

By generation 10,000, when mutation-selection balance is reached, simulations with the pleiotropic architecture generally maintain a higher average genetic correlation than those with linkage architectures, even when recombination is absent (linkage distance of 0cM between pairs of loci) (Figure 2). Variation in the mutation rate has the largest effect on the difference of genetic correlation between pleiotropic and fully linked non-pleiotropic loci, with much lower correlations as the mutation rate decreases from 10^−3^ to 10^−5^ (Figure 3). This reduction in genetic correlation mostly affected the non-pleiotropic pairs of loci for which a large drop in genetic correlation occurred between *µ* = 10^−3^ and *µ* = 10^−4^ (Figure 3). With lower mutation rates there is also a lower total genetic variance and lower genetic covariance. The higher genetic correlation obtained with pleiotropic loci was due to a lower total genetic variance when the mutation rate was high (*µ* = 10^−3^), but to a higher genetic covariance when mutation rate was low (*µ* = 10^−4^ or 10^−5^).

**Figure 2:**
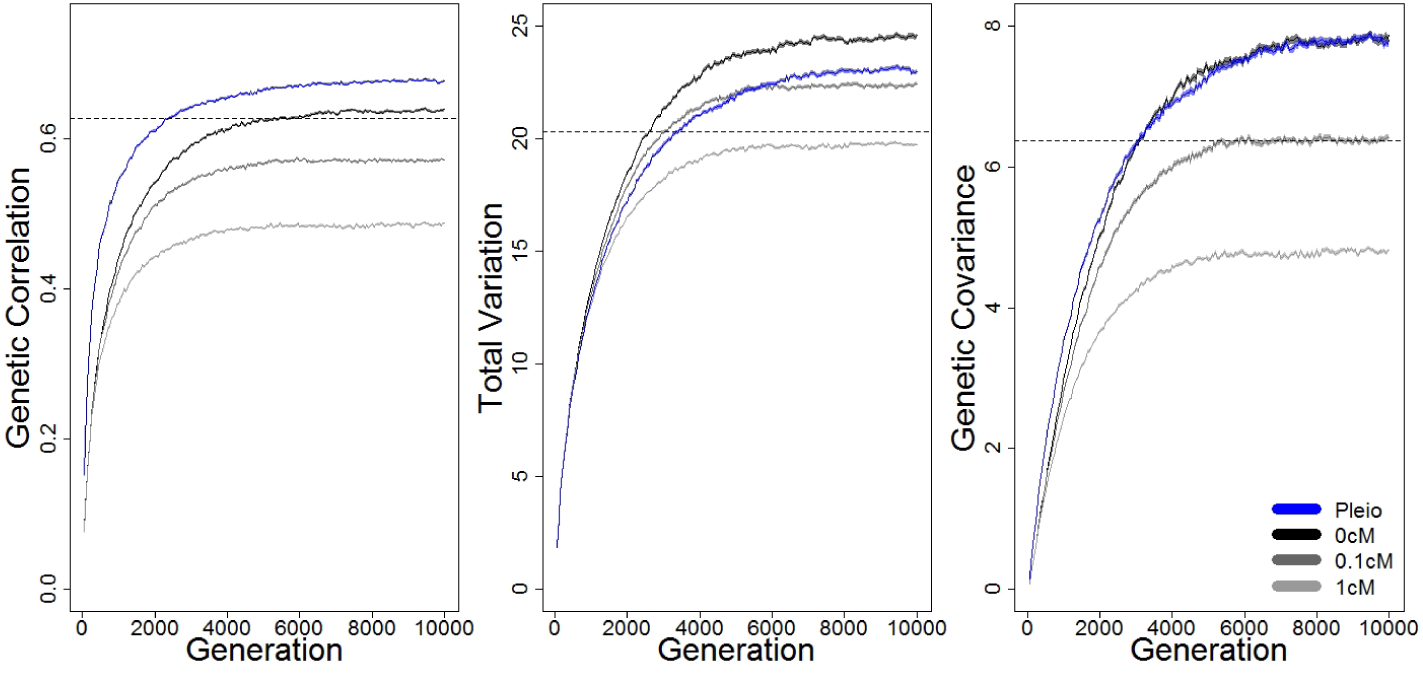
Average genetic correlation, total genetic variation and genetic covariation (and their standard deviations) over 10,000 generations reaching mutation-selection equilibrium for four different genetic architectures: pairs of linked loci affecting two different traits with 0, 0.1 or 1cM between loci, or pleiotropic loci affecting both traits. *N* = 5000, *ω*^2^ = 100, *ρ_ω_* = 0.9, *α*^2^ = 0.1, and *µ* = 0.001. Dashed line represents Lande’s 1984 expectations for completely linked loci (0 cm).

**Figure 3:**
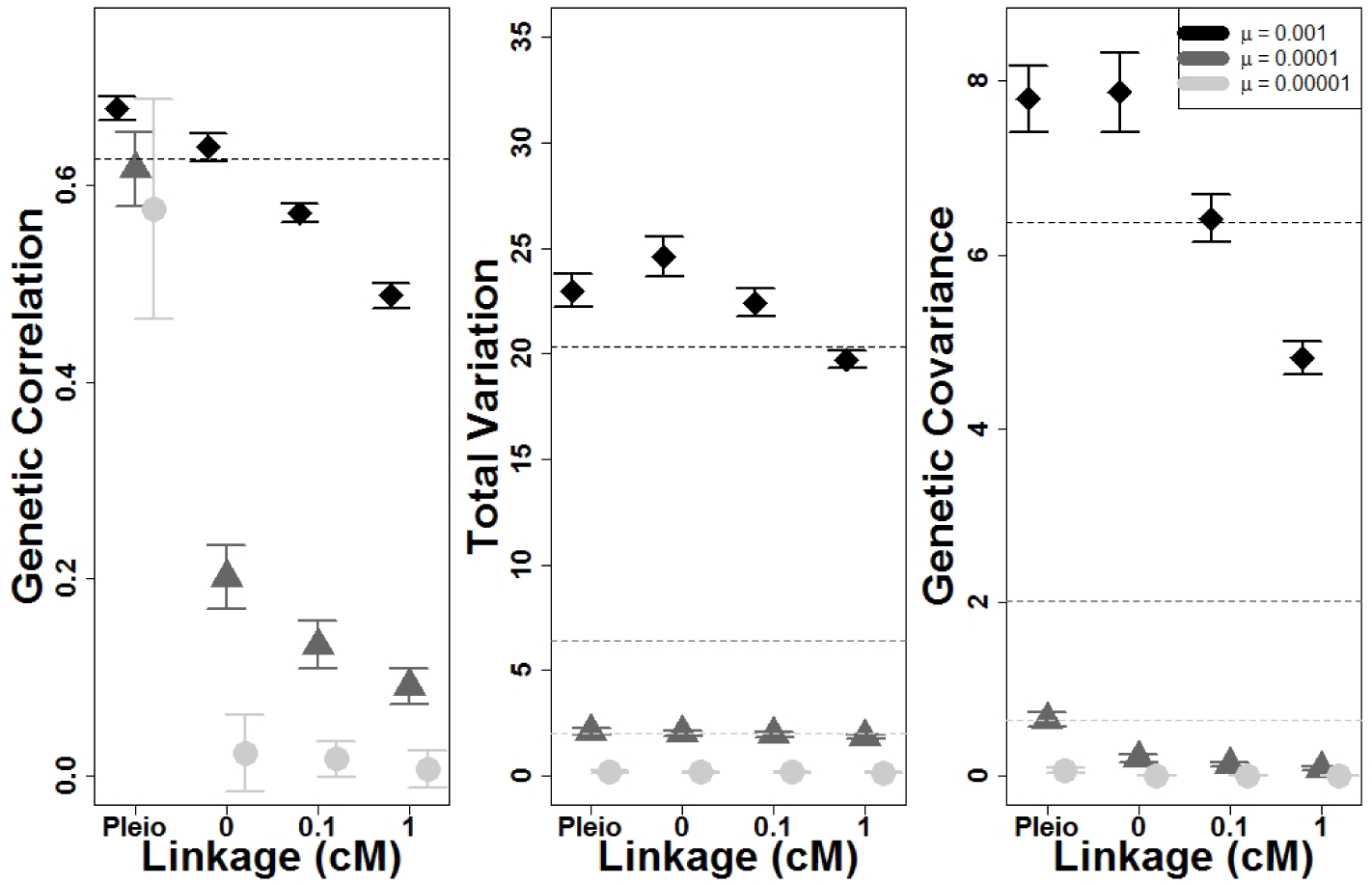
Effect of mutation rate (*µ*) on average genetic correlation, total variance and genetic covariance (and their standard deviations) after 10,000 generations of correlated, stabilizing selection for four different genetic architectures. *N* = 5000, *ω*^2^ = 100, *ρ_ω_* = 0.9, and *α*^2^ = 0.1. Dashed lines represents Lande’s 1984 expectations for completely linked loci (0 cM).

The genetic correlation between the traits decreases with reduction in all four factors tested (*µ*, *ρ_ω_*, *ω*^2^, and *α*^2^) and for all genetic architectures, with the coefficient of correlational selection (*ρ_ω_*) having the strongest effect (Figure 4), as expected from equation (3). However, changes in the strength of selection (*ω*^2^) and the mutational variance (*α*^2^) also affect the genetic correlation at equilibrium. We find that reducing the strength of selection (Figure 5) had a relatively smaller effect than reducing the mutational variance (Figure 6). A decrease in mutational variance leads to a decrease in genetic correlation by a similar amount regardless of genetic architecture (though loose linkage is affected the most). Populations with linkage architectures need both high mutation rates and high mutational variance to maintain strong genetic correlation, whereas the pleiotropic architecture just needs high mutational variance.

**Figure 4:**
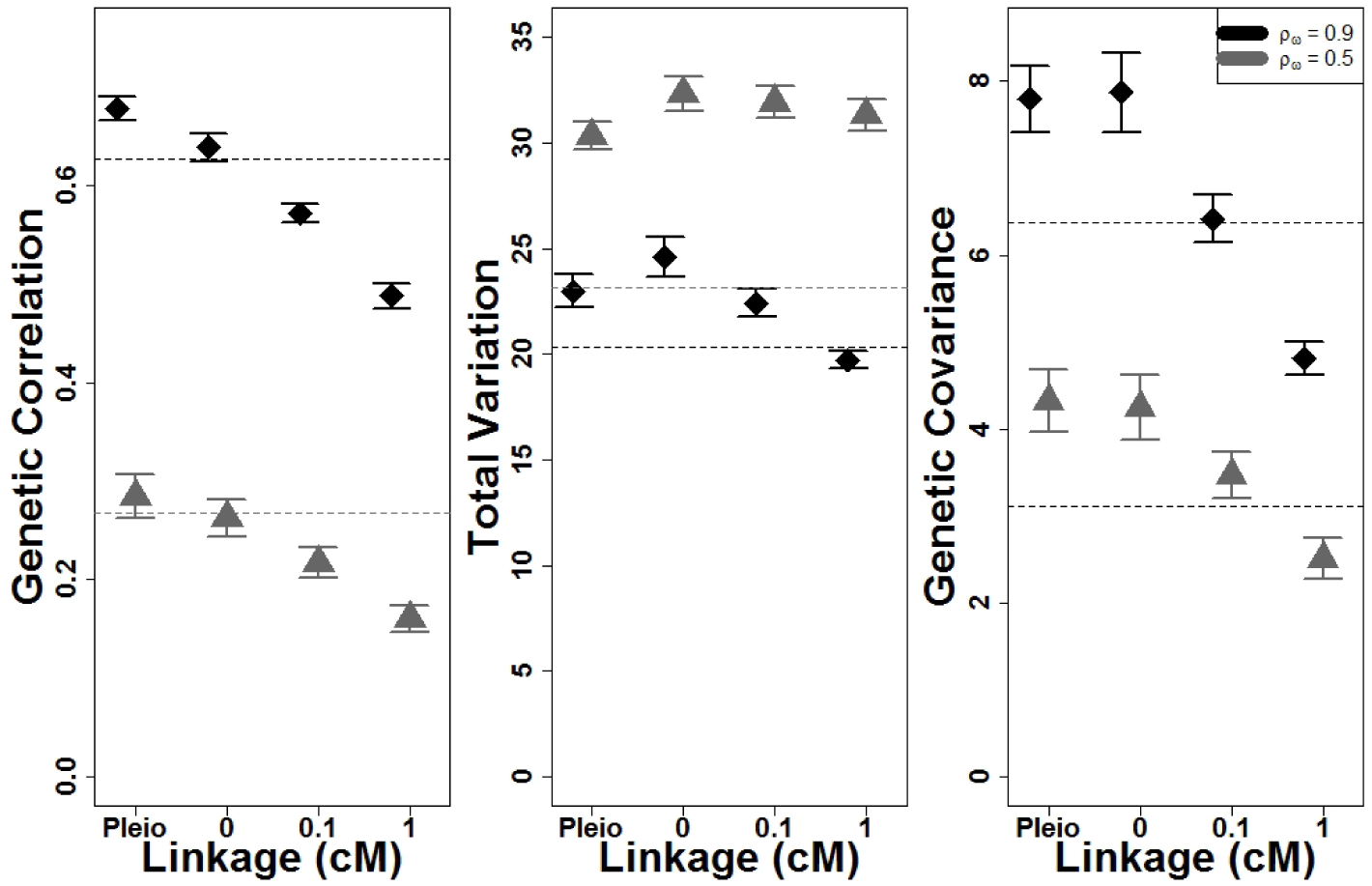
Effect of correlational selection (*ρ_ω_*) on average genetic correlation, total variance and genetic covariance (and their standard deviations) after 10,000 generations of correlated, stabilizing selection for four different genetic architectures. *N* = 5000, *ω*^2^ = 100, *α*^2^ = 0.1, and *µ* = 0.001. Dashed lines represents Lande’s 1984 expectations for completely linked loci (0 cM).

**Figure 5:**
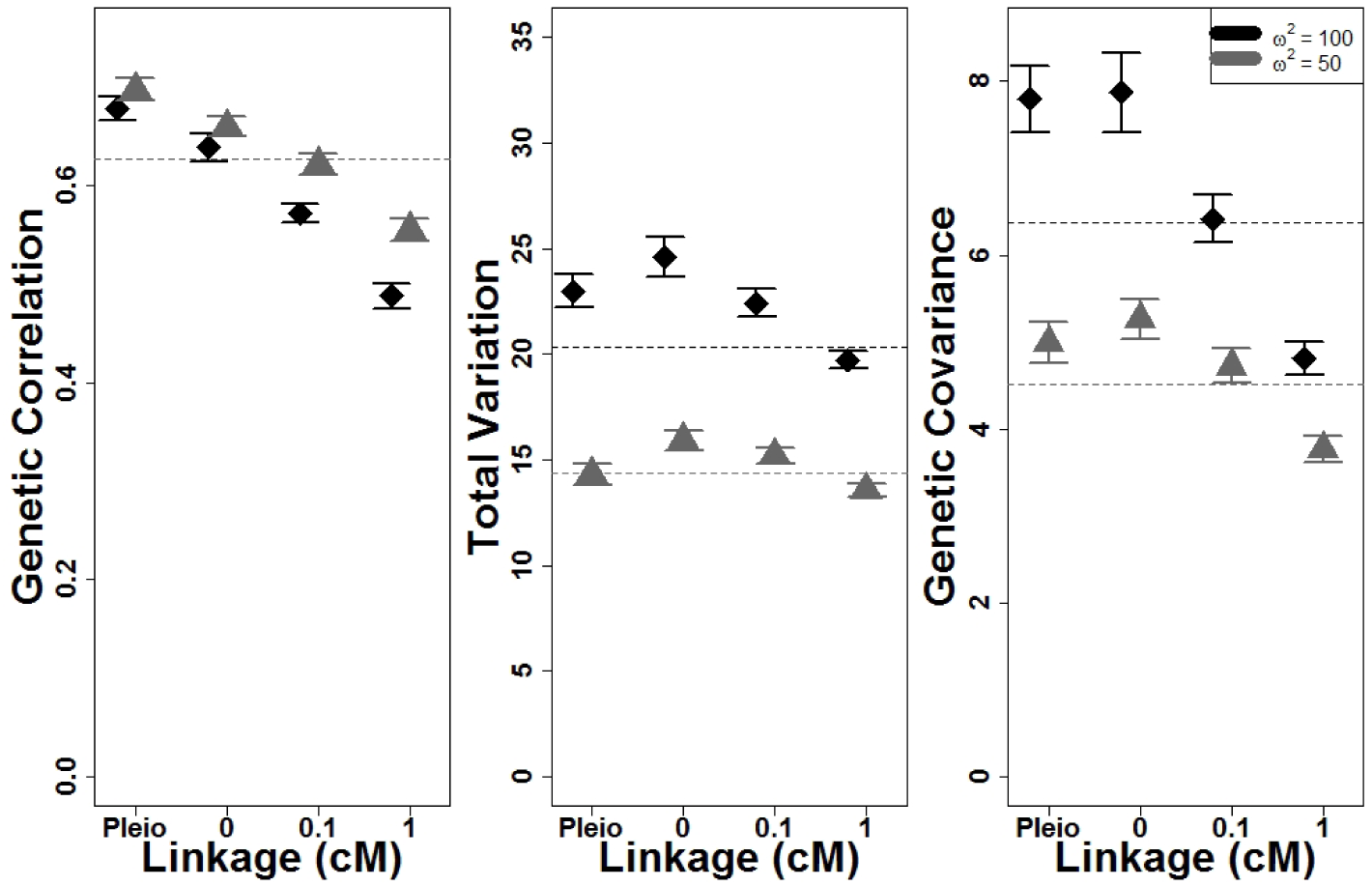
Effect of selection variance (*ω*^2^) on average genetic correlation, total variance and genetic covariance (and their standard deviations) after 10,000 generations of correlated, stabilizing selection for four different genetic architectures. *N* = 5000, *ρ_ω_* = 0.9, *α*^2^ = 0.1, and *µ* = 0.001. Dashed lines represents Lande’s 1984 expectations for completely linked loci (0 cM).

**Figure 6:**
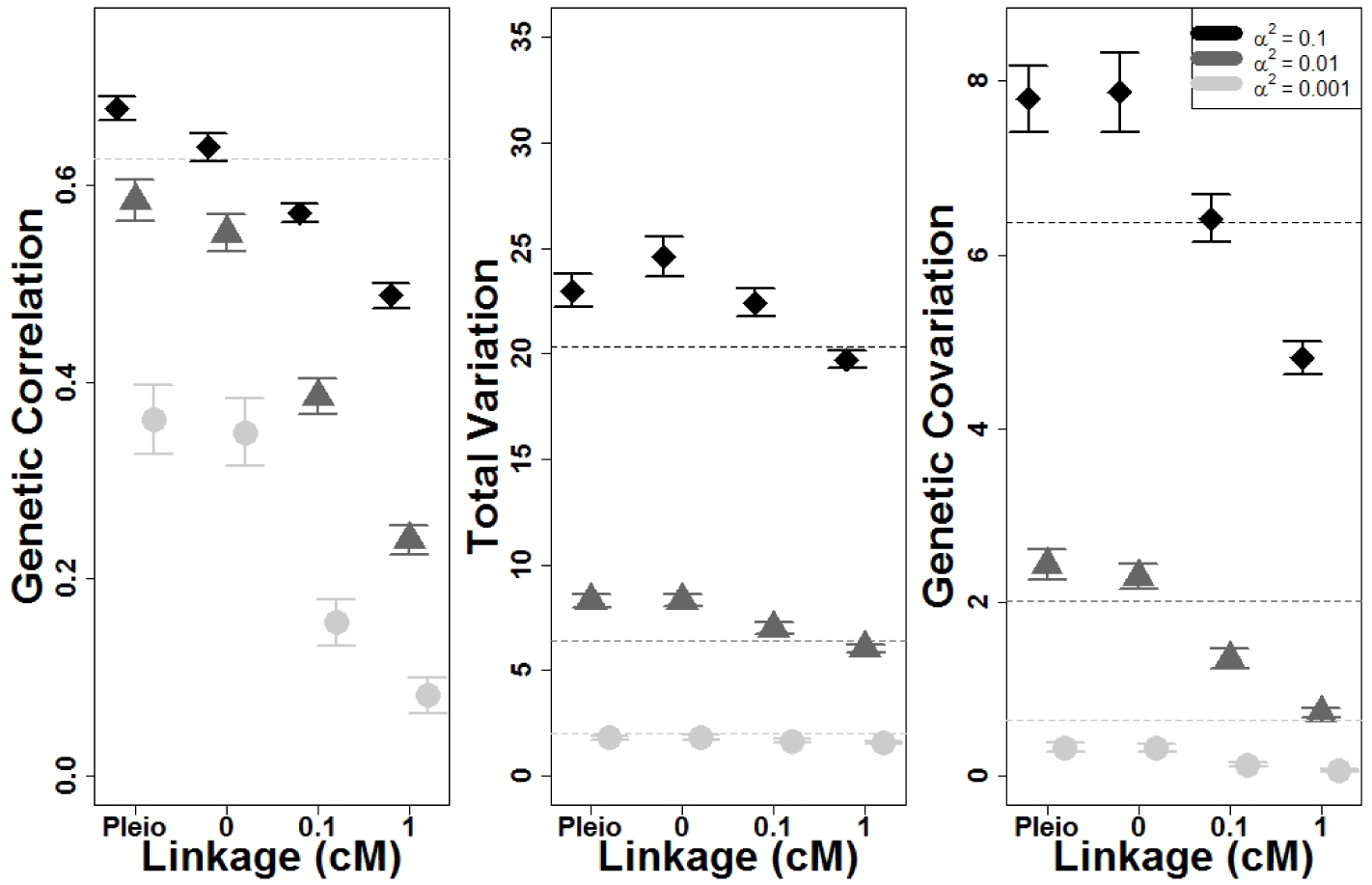
Effect of mutation variance (*α*^2^) on average genetic correlation, total variance and genetic covariance (and their standard deviations) after 10,000 generations of correlated, stabilizing selection for four different genetic architectures. *N* = 5000, *ω*^2^ = 100, *ρ_ω_* = 0.9, and *µ* = 0.001. Dashed lines represents Lande’s 1984 expectations for completely linked loci (0 cM).

In contrast to the correlation, the genetic covariance of the two traits was generally equal between pleiotropic and fully linked non-pleiotropic loci, and decreased as recombination increased within pairs of non-pleiotropic loci. The cause of the observed higher trait correlation obtained with pleiotropic loci was the lower genetic variance they maintain under stabilizing selection.

### Effects of migration on genetic correlation

A higher migration rate from a source population, whose traits are under correlational selection, leads to higher genetic correlations in the focal population regardless of the genetic architecture (Figure 7A). The effect of migration increases with tighter linkage and is highest with pleiotropic architecture. This effect on genetic correlation is still observed when there is no correlational selection on the traits in the source population, but to a largely reduced degree (Figure 7B).

**Figure 7:**
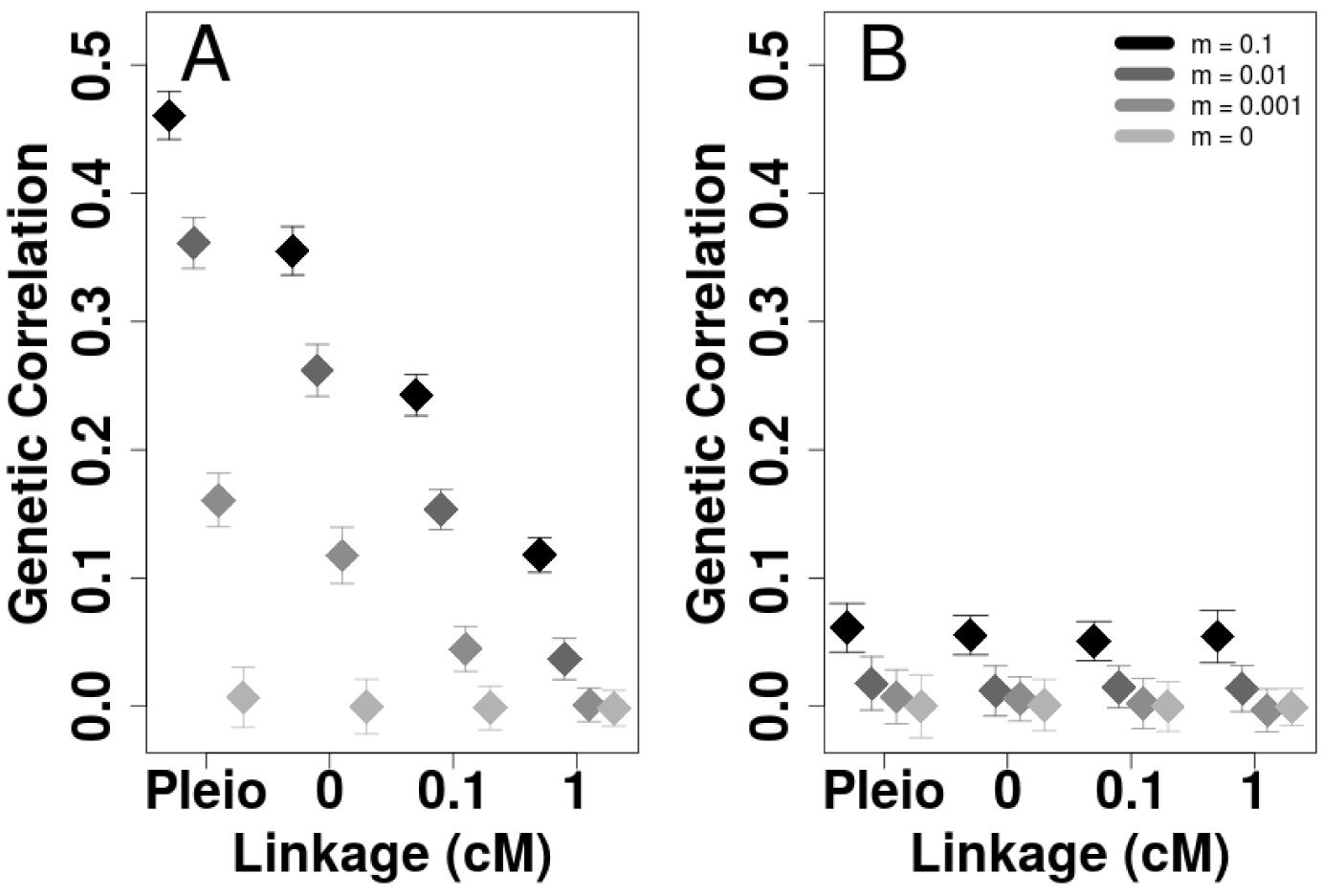
Average genetic correlations in the focal populations (and their standard deviations) after 10,000 generations of migration from a source population with different migration rates (*m*) for four different genetic architectures. A– Migration from a source population with correlational selection between traits (*ρ_ω_* = 0.9). B– Migration from a source population without correlational selection between traits (*ρ_ω_* = 0).

### Effects of linkage and pleiotropy on proportion of false positives/negatives and linkage disequilibrium in multi-trait GWASes

In simulations where there is linkage between SNPs and equivalent levels of genetic correlation between traits, the number and proportion of loci that are false positives (above FDR cutoff but no effect on trait) increase as linkage distance decreases between SNPs affecting different traits (shown in Figure 8 and Supplementary Figure S1). When genetic correlation is higher (due to stronger correlational selection), linkage distance has a greater impact on the proportion of false positives. Also, genetic correlation has a larger effect than linkage distance on the number of false positives. In simulations where SNPs are pleiotropic, genetic correlation due to correlational selection has little impact on the number and proportion of false negatives (below FDR cutoff but does affect the traits). Linkage disequilibrium between pairs of linked SNPs decreases as distance between SNPs increases regardless of genetic correlation (Figure 9 and Supplemental Table S1). Long-distance linkage disequilibrium between unlinked SNPs increases with the strength of correlational selection when the map distance within pairs of linked SNPs increases (when measured with *D′*, Supplemental Figure S2). In simulations where SNPs are pleiotropic, long-distance linkage disequilibrium does not seem to be affected by a change in genetic correlation. Finally, in simulations with neutral QTL, the false pleiotropic positive rate is 4.2e-5 (genetic correlation: *g*_*cor*_ = 0.2), 8.3e-5 (*g*_*cor*_ = 0.3), and 1.5e-4 (*g*_*cor*_ = 0.4) on average in the first 50kb window (within 0.05 cM of the causal QTL). No false pleiotropic positives were found at map distance above 0.15 cM for *g*_*cor*_ = 0.2 and 0.3, and above 0.25 cM for *g*_*cor*_ = 0.4.

**Figure 8:**
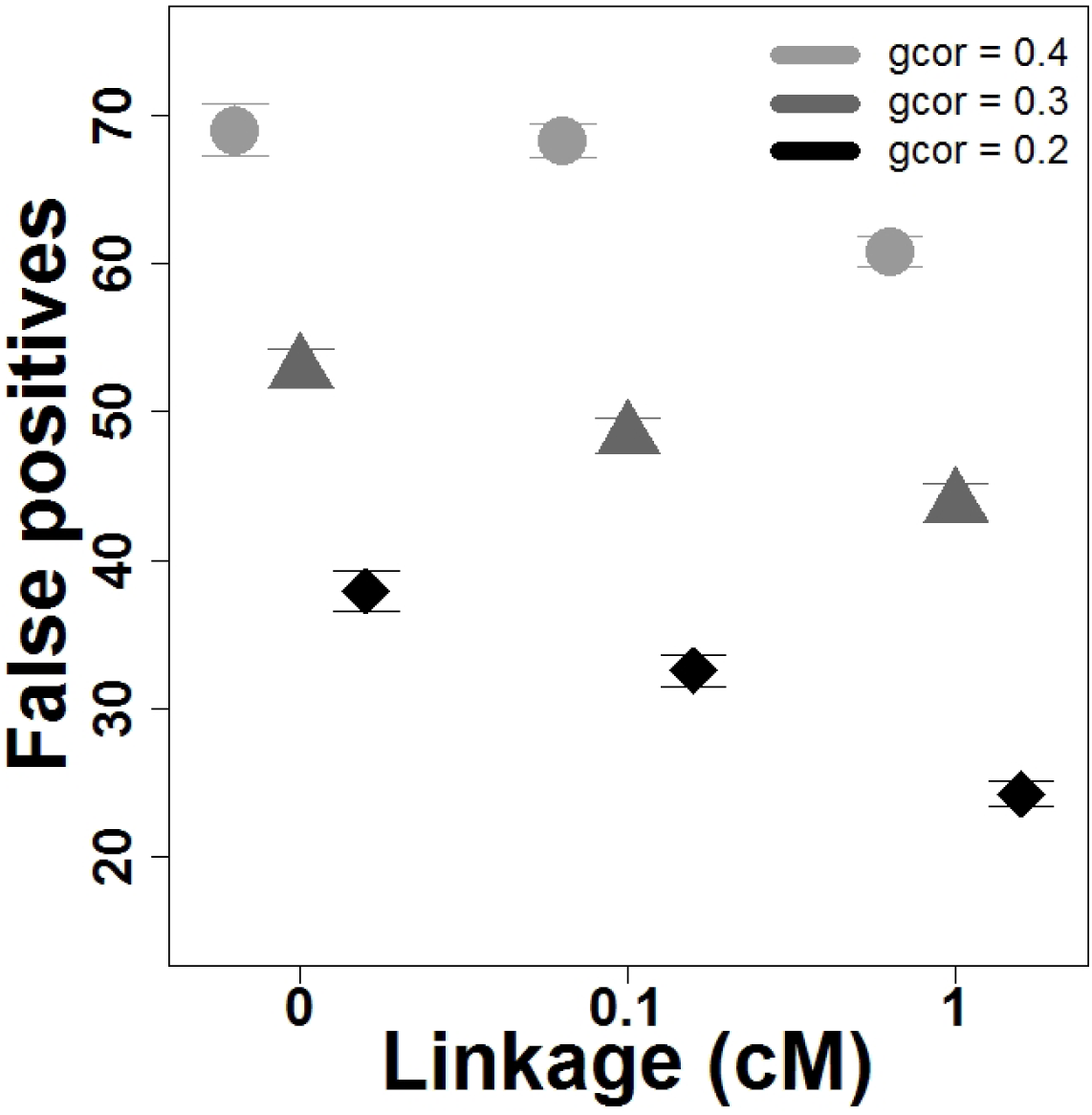
Average number of false positives from GWA analyses (and their standard deviations) for different linkage distances between paired loci and different genetic correlations (gcor). A locus was considered a false positive if associations between the locus’ genotypes and trait values, that the locus does not directly affect, are above the Benjamini-Hochberg FDR cutoffs (with a significance level of 0.05).

**Figure 9:**
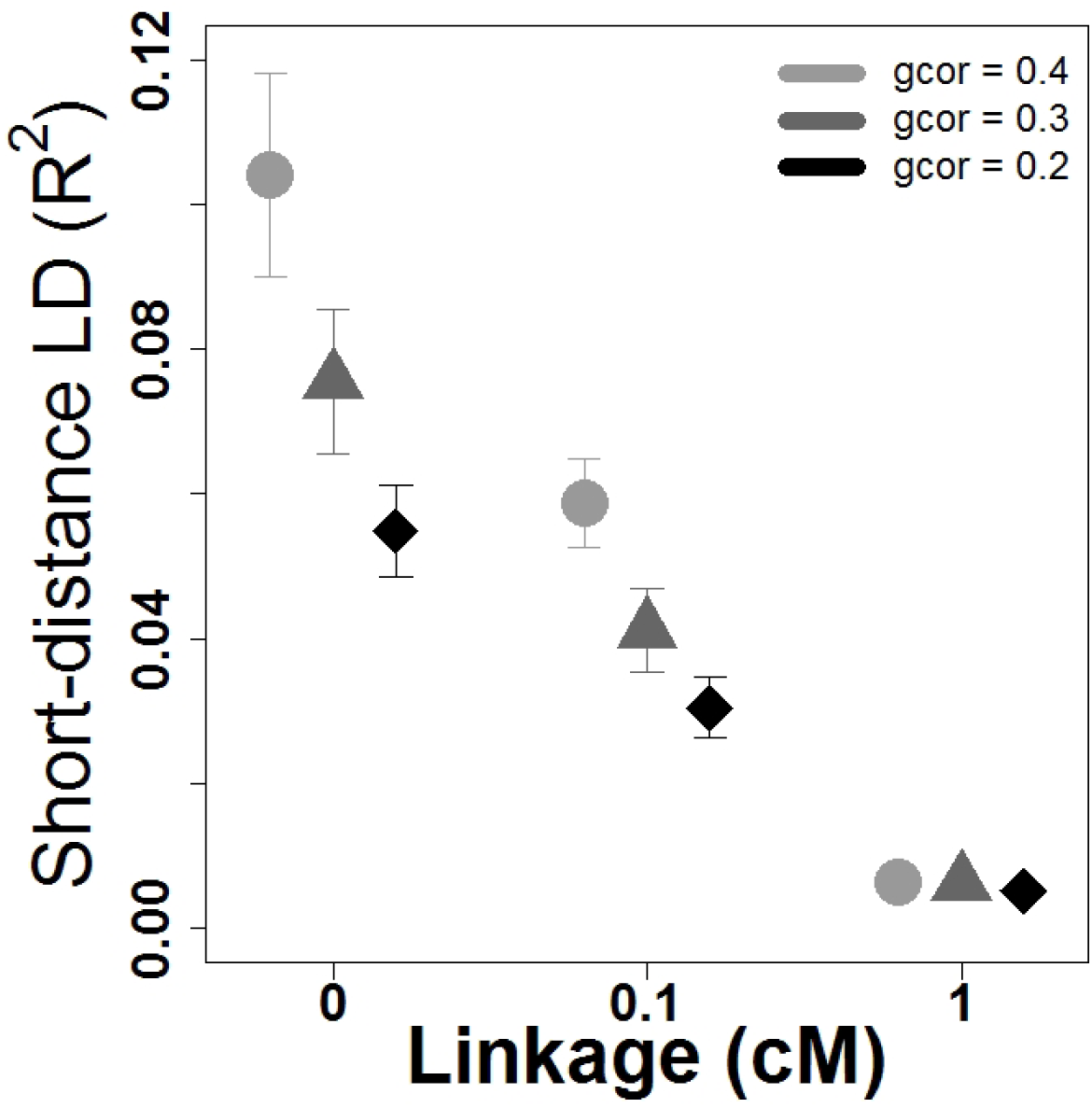
Average linkage disequilibrium (LD) between pairs of linked loci (and their standard deviations) for different linkage distances between paired loci and different genetic correlations (gcor).

## Discussion

### Pleiotropy and linkage are not the same

The main expectation under an assumption of weak selection and strong correlational selection is that populations with a genetic architecture consisting of unlinked pairs of two completely linked loci (0cM distance) should maintain similar equilibrium levels of genetic correlation as with a genetic architecture consisting of a lesser number of unlinked pleiotropic loci (Lande, 1984). Our results show that this is the case when there are half as many pleiotropic loci and mutation rates are relatively high. A high rate of mutation (10^−3^) allows for multiple mutations in both loci in a tightly linked pair to accumulate and maintain levels of genetic covariance near to that of mutations in a single pleiotropic locus, but empirical estimations of mutation rates from varied species like bacteria and humans suggests that *per-nucleotide* mutation rates are in the order of 10^−8^ to 10^−9^ (Nachman and Crowell, 2000; Ford et al., 2011; Keightley et al., 2015; Lindsay et al., 2019). If a polygenic locus consists of hundreds or thousands of nucleotides, as in the case of many quantitative trait loci (QTLs), then per-locus mutation rates may be as high as 10^−5^, but the larger the locus the higher the chance of recombination between within-locus variants that are contributing to genetic correlation. This leads us to believe that with empirically estimated levels of mutation and recombination, strong genetic correlation between traits are more likely to be maintained if there is an underlying pleiotropic architecture affecting them than will be maintained due to tight linkage. Consequently, GWASes that detect associations between multiple traits and single genetic variants are more likely to be detecting pleiotropic loci than linked loci. Also, previous theoretical models suggest that Lande’s (1984) equilibrium levels of genetic variation are not well approximated at low per-locus mutation rates (compared to the strength of selection), which was also true in our simulations (Supplemental Figure S3) (Turelli, 1984; Bürger, 2000).

We find that even under scenarios where pleiotropy and tight linkage maintain similar levels of genetic covariance, pleiotropic architectures have higher genetic correlations because they have lower total genetic variance. This can be explained by understanding the differential fitness effects of loci. Mutations that affect more than one trait are less likely to be beneficial (Orr, 1998; Otto, 2004). The distribution of fitness effects of pleiotropic mutations is shifted towards more negative average values as the number of traits affected increases (Martin and Lenormand, 2006; Chevin et al., 2010). Hence, pleiotropic architectures that affect more traits have less positive mutational effects on fitness and maintain a lower equilibrium genetic variation when compared to linked architectures (Turelli, 1985). It has been suggested that this might be overcome in more complex organisms with a greater number of traits by modularization of the effects of different pleiotropic genes to separate sets of traits and decrease the pleiotropic degree of the mutations but theoretical models have shown mixed results (Baatz and Wagner, 1997; Hansen, 2003; Welch et al., 2003; Martin and Lenormand, 2006; Chevin et al., 2010; Wagner and Zhang, 2011).

When correlational selection on the traits is strong in the simulations with linked architectures, the equilibrium genetic correlation is dependent on the recombination rates between loci within linkage groups. Tightly linked loci can maintain higher levels of genetic correlation from a build-up of positive linkage disequilibrium than loosely linked loci. This matches the analytical predictions put forth in Lande (1984) under the assumption of weak stabilizing selection, strong correlational selection, and loose linkage between loci affecting the same trait.

### The impact of pleiotropy and linkage maintaining different genetic correlations in association studies

When methods like GWA analyses are employed to detect shared genetic influences (pleiotropy or linkage) on multiple traits of interest, they are dependent upon detecting combinations of effect sizes of genetic variants associated with those traits (Hill and Zhang, 2012b,a; Chung et al., 2014; Visscher and Yang, 2016). The success or failure of this endeavor is directly connected to the ability to detect loci with associations to each trait and the strength of genetic correlation between traits (Wei et al., 2014; Pickrell et al., 2016; Chesmore et al., 2018; Verbanck et al., 2018). Our results show that (tight) linkage between loci affecting different but correlated traits will lead to “many” false positives. We also show that false positive rates are only marginally affected by linked but non-causative loci. As expected, false positive rates decrease with the map distance to the causative loci and the correlation between the traits (Siegmund and Yakir, 2007). Therefore, GWASes will not be able to empirically distinguish between pleiotropy and linkage when loci affect genetically correlated traits. The proportion of genes associated with two or more phenotypes in the GWAS catalog has increased to around 40% in the last decade (Welter et al., 2013; Pickrell et al., 2016). But it is difficult to determine if this is truly representative of the prevalence of pleiotropy because QTLs are often mapped to loci that can encompass thousands of nucleotides (and more than one gene) and informative SNPs with significant effect sizes are assigned to the closest genes with annotated phenotypes (Chesmore et al., 2018; Liu et al., 2019; Cai et al., 2020). Conflating inter-genic SNPs with nearby pleiotropic genes (or loci) can distort the prevalence of pleiotropy and reduce the ability to distinguish pleiotropy from physical linkage (Dudley et al., 2005; Gianola et al., 2015). Finding the true false positive rate in GWA studies due to linkage is difficult because it is almost never known whether the source of genetic correlations between traits is linked loci or not, even when fine-scale sequences are available (for the reasons mentioned above and because of the way pleiotropy is erroneously defined in GWA studies) (Platt et al., 2010). Watanabe et al. (2018) attempted to break down this issue in a meta-analysis of 558 GWASes by looking at the proportion of genomic loci, genes, and SNPs associated with multiple traits, which may provide a clearer picture of the prevalence of pleiotropic causal variants. They found that 93.3% of loci, 81.0% of genes, and 60.2% of SNPs, were associated with more than one trait. This may seem to provide a better estimate of pleiotropic levels, except that in this study SNPs that were associated with more than one trait could still have been the result of linkage disequilibrium. A point that was brought up by the authors.

On the other hand, we observed very few false negatives in pleiotropic loci (regardless of genetic correlation) because we “sampled” the entire population and therefore had the power to find significant associations with (almost) all causal loci. Had we taken smaller samples of our population to perform the GWA analysis, we would have found a greater number of false negatives. The salient consequence is that study design, threshold levels, and genetic correlations between traits will all affect detection of genetic variants, whether the variants are causal themselves or linked to causal variants (Wagner and Zhang, 2011; Hill and Zhang, 2012a). Also, the number of pleiotropic effects a locus has may be under-represented by significance levels in association studies (Hill and Zhang, 2012b). Wagner and Zhang 2011 go a step further to suggest that number or proportions of traits affected may not be as meaningful as describing the distributions of pleiotropic effect sizes on traits.

### There is a difference between pleiotropy and linkage at the nucleotide level

Transgenic experiments may differentiate pleiotropy from linkage at the gene level (Mills et al., 2014), but at the nucleotide level does the distinction between two linked loci and one pleiotropic locus go away? There is evidence that even in the same gene, adjacent polymorphisms affecting different traits in *Drosophila* can be in linkage equilibrium due to fine-scale recombination (Carbone et al., 2006; Flint and Mackay, 2009). But imagine a case where a mutation in a single base-pair has an effect on one trait and a mutation in the base-pair right next to the first base-pair has an effect on a second trait. Now imagine a second case where a mutation in a single base-pair has an effect on two traits. There still seems to be a distinction between these two cases because the probability of a change in both traits in the first case is the mutation rate squared compared to the second case where the probability of a change in both traits is just the mutation rate. Depending on the per-locus mutation rate this difference can be quite large (e.g. 10^−8^ versus 10^−16^). Even in this extreme case, there may indeed still be a gray area in the distinction between pleiotropy and linkage at a mutational level. Mutations may affect the pleiotropic degree (e.g. like enzyme specificity) of a protein-coding gene and the degree to which the gene maintains multi-functionality may itself evolve (Guillaume and Otto, 2012). If there is correlational selection between the catalytic functions of an enzyme, then some pleiotropic mutations that affect more than one catalytic ability will be favoured, and genetic correlations will increase. With this in mind, it makes more sense from a theoretical and functional standpoint to refer to pleiotropy at the nucleotide level (or at the unit of a mutation), than at the gene or larger locus level (but this may depend on the questions of interest (Rockman, 2012; Rausher and Delph, 2015)).

### Other factors

Even in the absence of correlational selection it is possible to maintain genetic correlation through continued migration from a source population. High migration brings individuals whose combination of alleles will expand focal population variation in the direction of the source population. This corroborates previous results that showed that slow introgression of allelic combinations into a population can affect the genetic variance-covariance structure of that population (Guillaume and Whitlock, 2007). Whether genetic covariance will be maintained in real populations depends on the nature of correlational selection on traits in the population of interest, since migration can reduce local fitness (i.e. migration load) if allele combinations are not favoured by selection or increase it if they are (Nosil et al., 2006; Bolnick and Otto, 2013). Migration into a population will also affect false positive rates since immigrating allele combinations will be in LD from the source population and will therefore increase the proportion of certain genotypes, even if there is no strong trait correlation in the source population. Although not investigated in this study, a structured population and/or a continual system of inbreeding in a population where there is correlational selection between polygenic traits can result in increased genetic covariation caused by larger LD Lande (1984), which can in turn increase false positive proportions.

## Conclusion

Pleiotropic loci maintain stronger genetic correlations between traits than linked loci affecting different traits even when no recombination occurs between the loci, and especially in the magnitude of empirically estimated mutation rates. Previous models of the maintenance of genetic covariation at mutation-selection equilibrium describe genetic covariation as a function of the product of mutation rate and variance. These models provide similar expectations for pleiotropic and tight linkage architectures. The discrepancy occurs because of the contingency of mutational covariance input on the occurrence of mutations (and hence mutation rate). Without high mutation rates, the ability to create genetic covariance between linked loci is highly diminished because the combined likelihood of mutations in each linked loci with both mutational effects in the same direction is low. This result will have implications in the type of underlying architecture we expect to find in multi-trait association studies. On the one hand, tighter linkage between causal loci and detected loci maintains higher genetic correlations, leading to a greater proportion of false positives in pleiotropy tests. More importantly, on the other hand variants are more likely to have pleiotropic effects on traits than linked effects, when they are found to be associated with strongly correlated traits.

## Acknowledgments

This manuscript benefited from the comments and constructive criticism provided by Kathleen Lotterhos, Pär Ingvarsson, and an anonymous reviewer at PCI Evolutionary Biology. Version 3 of this preprint has been peer-reviewed and recommended by Peer Community In Evolutionary Biology (https://doi.org/10.24072/pci.evolbiol.100087). J.C. and F.G. were supported by the Swiss National Science Foundation, grant PP00P3 144846/1 to F.G.

## Author Contributions

J.C. performed software modification for model implementation and acquisition of data, as well as drafting of manuscript. J.C. and F.G. performed study conception and design, analysis and interpretation of data, and critical revision of manuscript.

## Data Archival

The data for this study will be made available online through Zenodo online repository at https://zenodo.org/record/3370185#collapseTwo and code for simulations can be found https://sourceforge.net/projects/nemo2/files/PublicationsCode/ChebibGuillaume-PleiotropyOrLinkage-2019/

## Supplemental

**Figure S1:**
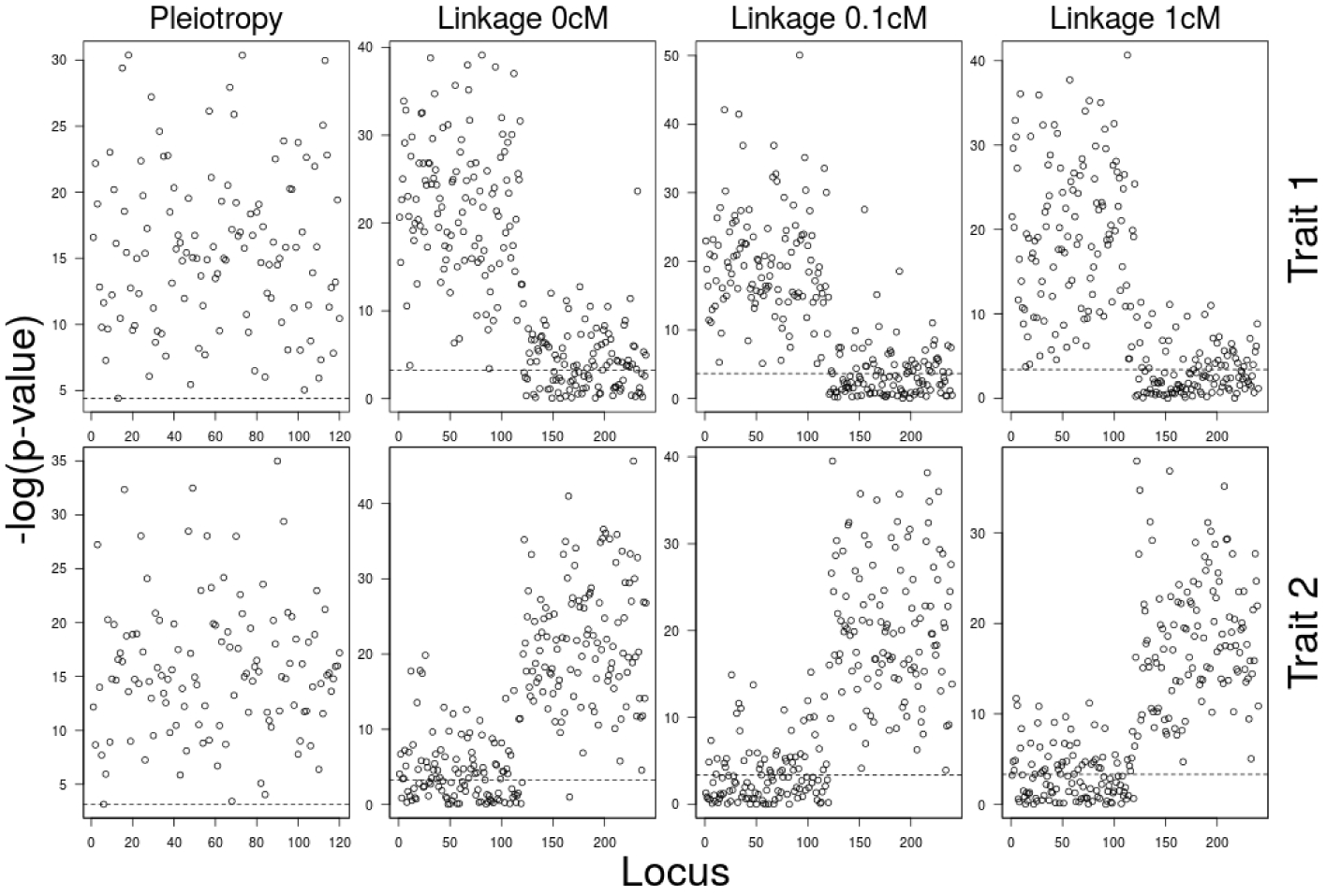
GWA analysis: -log(p-values of slope of regression of trait values on genotypes) from one set of example simulations. In the case of linkage architectures, the first 120 loci only affected trait 1 and the next 120 loci only affected trait 2. The order of the loci are sorted for visualization purposes whereby linked pairs are separated by the trait they affect (e.g. loci 1 and 121 in the figure are a linked pair). In the case of the pleiotropic architecture, all 120 loci affected both traits. The average genetic correlation of ≈ 0.3 was observed by adjusting the correlational selection levels to 0.88, 0.89, 0.93, and 0.965 for pleiotropy, linkage 0cM, linkage 0.1cM, and linkage 1cM, respectively. Dashed lines represent the Benjamini-Hochberg FDR cutoffs for a significance level of 0.05.

**Figure S2:**
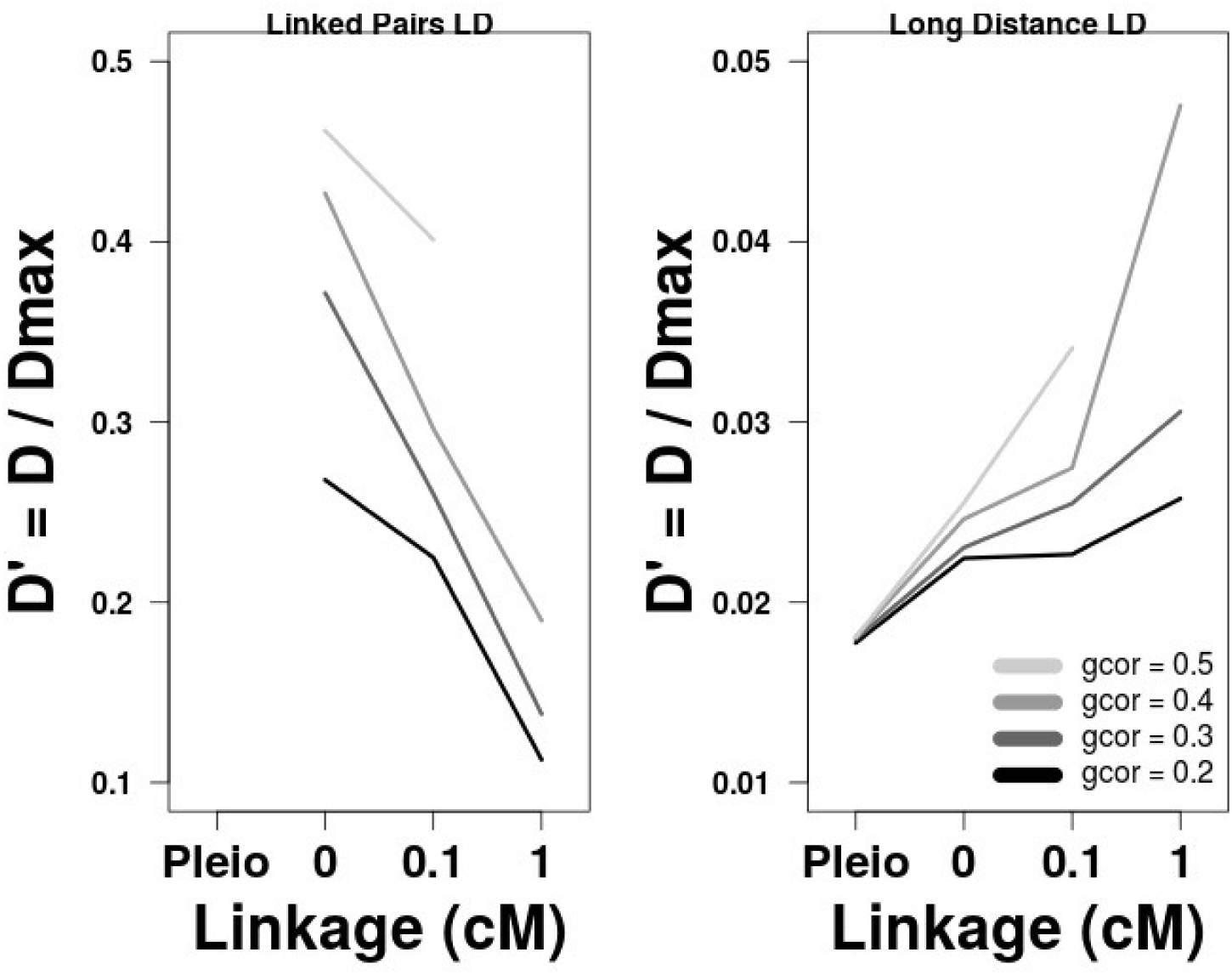
Average linkage disequilibrium (measured by *D′*) between linked pairs (left panel) and between unlinked pairs (right panels) for different genetic correlations and genetic architectures. N.B. No linked pairs existed between pleiotropic loci.

**Figure S3:**
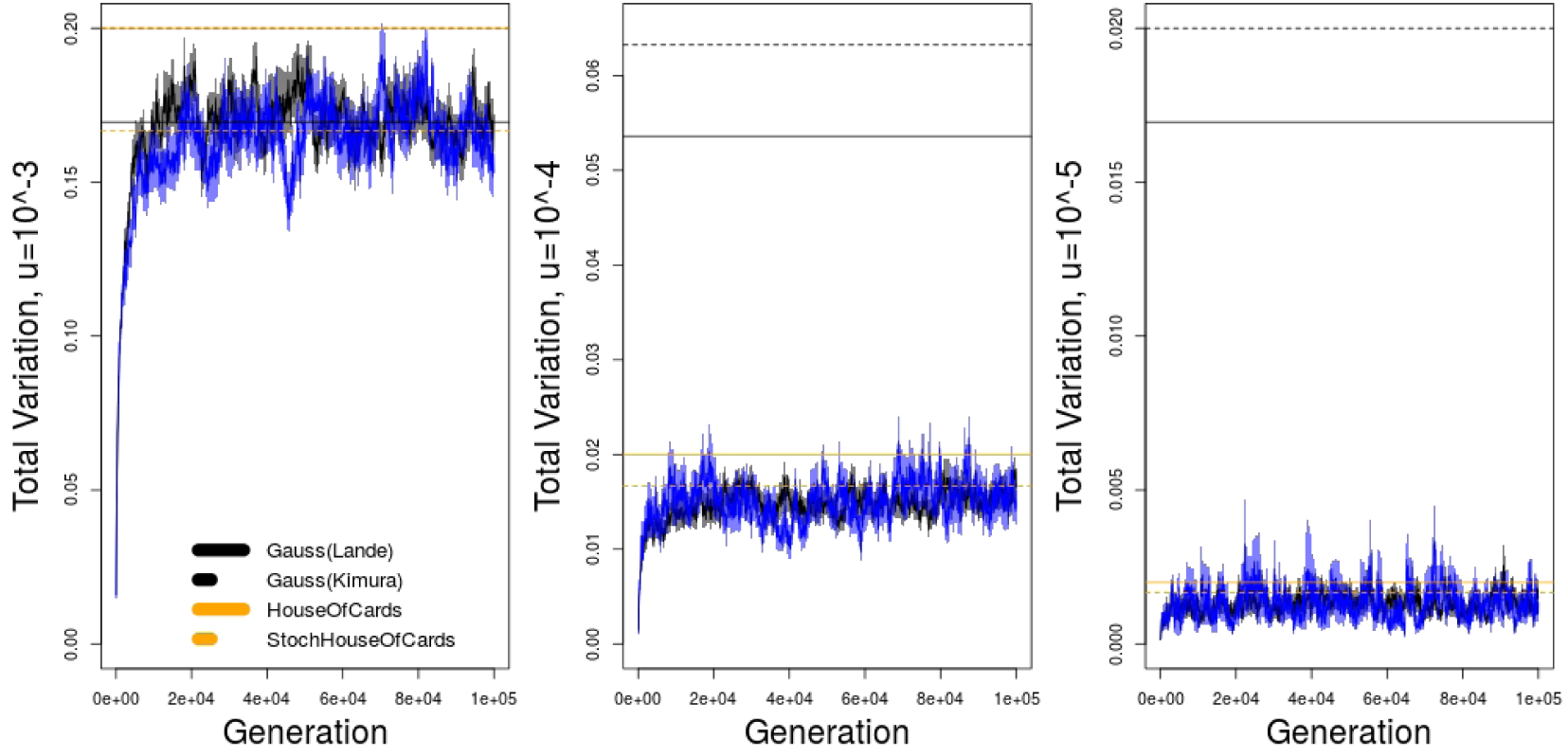
Average genetic variances for different mutation rates and genetic architectures, with either one pleiotropic locus or two completely linked loci, compared against theoretical expectations from several models (Bürger, 2000).

**Table S1:** Results of GWA analyses for different architectures with average false negatives (Type II errors) for pleiotropic architectures and false positives (Type I errors) for linkage architectures, as well as linkage disequilibrium (LD) measurement averages for short-distance (physically linked loci) and long-distance (unlinked loci) comparisons. The genetic architectures in the bottom half of the table have higher genetic correlations than the top half (created by adjusting correlational selection) to compare the differences at different genetic correlation.

| Genetic Architecture | Genetic Cor (SE) | Type I/II Error % | D' short | D' long | $R^2$ short | $R^2$ long |
| --- | --- | --- | --- | --- | --- | --- |
| Pleiotropy | 0.308 (0.0046) | 0.35% | NA | 0.018 | NA | 0.00027 |
| Linkage (0cM) | 0.300 (0.0055) | 22.06% | 0.37 | 0.023 | 0.089 | 0.00026 |
| Linkage (0.1cM) | 0.300 (0.0045) | 20.17% | 0.26 | 0.025 | 0.047 | 0.00027 |
| Linkage (1cM) | 0.308 (0.0035) | 18.28% | 0.13 | 0.030 | 0.007 | 0.00027 |
| Pleiotropy | 0.407 (0.0048) | 0.32% | NA | 0.018 | NA | 0.00027 |
| Linkage (0cM) | 0.398 (0.0074) | 28.76% | 0.43 | 0.025 | 0.107 | 0.00027 |
| Linkage (0.1cM) | 0.408 (0.0035) | 28.46% | 0.30 | 0.027 | 0.050 | 0.00027 |
| Linkage (1cM) | 0.404 (0.0029) | 25.34% | 0.19 | 0.048 | 0.006 | 0.00027 |

